# Association between COVID-19 Disease Severity and T Cell Receptor Repertoire

**DOI:** 10.1101/2024.11.26.625333

**Authors:** Xinyi Chen, Yapeng Su, Chloe Dai, Jason D. Goldman, James R. Heath, Si Liu, Wei Sun

## Abstract

During the COVID-19 pandemic, while most infected individuals experienced mild to moderate symptoms, a significant subset developed severe illness. A clinical test distinguishing between mild and severe cases could inform effective treatment strategies. Toward the latter stages of the pandemic, it became evident that vaccination or prior infection cannot entirely prevent reinfection. However, they are crucial in reducing the risk of severe disease by inducing T-cell memory. T cell receptors (TCRs), which can be obtained from human blood, serve as valuable biomarkers for monitoring T cell responses to SARS-CoV-2 infection. In this study, we investigated the associations between TCR metrics and COVID-19 severity and found significant associations. Furthermore, such associations could depend on the subset of TCRs used (e.g., TCRs from CD8+ T or CD4+ T cells) and when the TCRs were collected.

## Introduction

T cells are crucial parts of the human immune system and play a central role in the recognition and combating of infections. Infected cells present fragments of foreign proteins, which are identified by T cells through their T Cell Receptors (TCRs). Each T cell typically expresses a distinct TCR, and the collection of millions of distinct TCRs within an individual is known as a TCR repertoire [1]. TCR repertoires are valuable resources for developing biomarkers for a range of immune-related diseases, including infectious diseases [2, 3], autoimmune conditions [4], cancer [5], and Alzheimer’s disease [6].

Several unique features of TCRs suggest that they may meet unmet clinical needs [2]. First, a TCR can have high sensitivity and specificity to detect a small amount of antigen. Second, TCRs can retain a memory of infections for many years, providing a durable record of immune history. Finally, TCRs can be conveniently collected through a simple blood draw. Despite these advantages, TCRs remained an underutilized resource until recent advances in sequencing technologies made it possible to accurately analyze large numbers of TCRs [7,8].

Pioneering studies have demonstrated that TCRs associated with virus infection (e.g., cytomegalovirus (CMV) [2] or SARS-CoV-2 [3]) can be identified through case-control studies. These infection-specific TCRs can serve as markers of current or past infections. Several studies have revealed that vaccination and natural infection induce T cell memory, a critical factor in protecting against severe diseases [9–11]. Motivated by these studies, our aim is to investigate the associations between TCRs and COVID-19 severity.

We hypothesize that a robust T cell response to SARS-CoV-2 during the early stages of COVID-19 is associated with milder disease outcomes. To test this hypothesis, we quantify T cell responses using the abundance of SARS-CoV-2-associated TCRs (SC2-TCRs for short), and associate them with COVID-19 severity. In this paper, all SC2-TCRs were obtained from a dataset [3,12] generated through MIRA (Multiplex Identification of Antigen-Specific T-Cell Receptors) technology, a technology that allows for the simultaneous identification of T cells specific to a large number of antigens. The MIRA dataset includes more than 140,000 TCRs that can be divided into two groups: (1) HLA-I TCRs, or CD8+ TCRs, originating from CD8+ T cells reacting to peptides presented by HLA-I proteins, and (2) HLA-II TCRs, or CD4+ TCRs, derived from CD4+ T cells reacting to peptides presented by HLA-II proteins. We analyze the TCR data reported by Su et al. [13] and Delmonte et al. [14], which were collected from COVID-19 patients with varying disease severities and at different time points.

Our findings reveal that higher abundance of CD8+ SC2-TCRs at early time points is associated with milder COVID-19. Interestingly, at later times, a higher abundance of CD4 + SC2-TCRs is associated with more severe diseases, suggesting a dynamic, time-dependent relationship between SC2-TCR profiles and disease severity. These results highlight the potential of TCR-based predictors as tools to predict disease progression, which could improve disease management and inform therapeutic strategies.

## Methods

### MIRA data processing

The MIRA dataset consists of 148,301 experimentally predicted SARS-CoV-2 peptide-associated TCR beta-chains [3, 12]. These TCRs were collected from PBMC (Peripheral Blood Mononuclear Cell) or T cell samples from 144 individuals, including 105 individuals with SARS-CoV-2 exposure (COVID set) and 39 individuals without documented SARS-CoV-2 exposure (healthy set) (Figure 1(A)). Previous studies have shown that SARS-CoV-2–specific TCRs can be detected in individuals without SARS-CoV-2 exposure because these TCRs cross-react with other antigens, such as common cold coronavirus [15], commensal bacteria [16], or cytomegalovirus (CMV) [17]. Therefore, the MIRA TCRs from the healthy set are not necessarily less reliable or informative than those from the COVID set.

**Figure 1.**
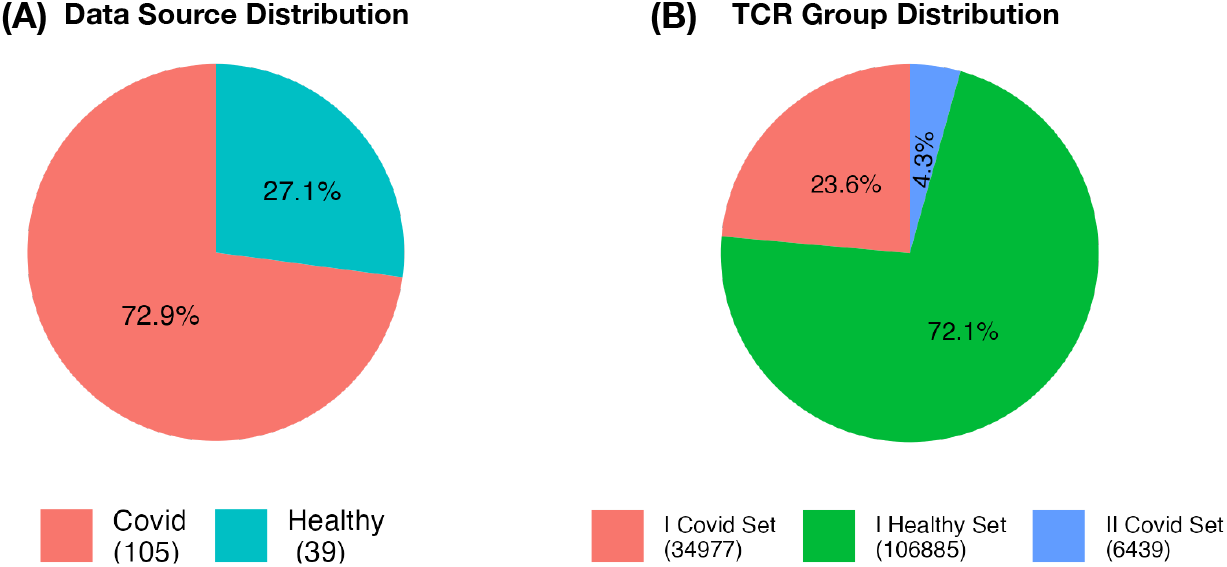
(A) Distribution of MIRA TCRs from 105 COVID-19 patients or 39 healthy individuals. (B) Distributions of MIRA TCRs in three groups.

We categorize the MIRA TCRs into three groups: CD8+ TCRs from the COVID set, CD8+ TCRs from the healthy set, and CD4+ TCRs, which are exclusively from the COVID set. Since CD8+ TCRs and CD4+ TCRs recognize antigens presented by HLA class I and HLA class II molecules, respectively, we also refer to them as (HLA class) I-COVID set, I-Healthy set, and II-COVID set, respectively (Figure 1(B)). This classification allows us to examine the associations between COVID-19 severity and the abundance of TCRs within each group. In the Results section, we demonstrate that CD8+ TCRs and CD4+ TCRs exhibit distinct association patterns with COVID-19 severity. However, we observe no significant differences between the I-COVID set and I-Healthy set in terms of their relationship with disease severity.

### Definition of TCR metrics

A TCR consists of an alpha chain and a beta chain, with the beta chain exhibiting greater sequence diversity. In a healthy individual, approximately 10^7^ unique TCR beta chains [2] are expressed from an effective pool of ∼ 10^14^ possible beta chains [18]. Due to such extreme sequence diversity of TCRs, all T cells sharing the same TCR must arise from the clonal expansion of a single T cell. This unique property allows TCRs to serve as identifiers for T cell clones.

The collection of all the TCRs from an individual is referred to as a TCR repertoire. To evaluate the proportion of SC2-TCRs within a TCR repertoire, we use two metrics: richness and abundance. A unique TCR is defined by its V gene, J gene, and CDR3 sequence. Denote the total number of TCR clones (i.e., the number of unique TCRs) of the *i*-th individual by *n*_*i*_, and the number of TCRs within each clone by *x*_*ij*_ for 1 ≤ *j* ≤ *n*_*i*_. Without loss of generality, assume the first *k*_*i*_ of the *n*_*i*_ unique TCRs are SC2-TCRs. Then we define the richness (*r*_*i*_) and abundance (*a*_*i*_) of MIRA TCRs as

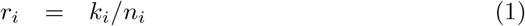

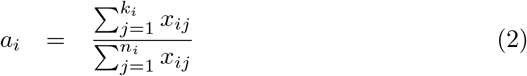

### Single cell and bulk TCR data from Su et al

Su et al. [13] reported TCR data from 265 COVID-19 samples collected from 139 COVID-19 patients varies from different time points. The samples were obtained from five hospitals of Swedish Medical Center and affiliated clinics in the Puget Sound region near Seattle, WA, before September 2020. Each sample was collected at a time point within 1-2 weeks after the onset of symptoms and consistently categorized into two sub-time points: early (T1) and later (T2). In this paper, we focused on 121 patients with TCR data available at the early (T1) time point, since we hope to use the TCR metric at the early time point to predict COVID-19 severity. Among the 121 patients, the sample sizes for males and females were relatively balanced, with 67 females and 54 males, and the median age was around 58. Disease severity were quantified using WHO Ordinal Scale for Clinical Improvement score (WOS) [19]: mild [WOS = 1–2, not hospitalized], moderate [WOS = 3–4, hospitalized, with or without oxygen], and severe [WOS = 5–7, hospitalized, with ventilation and treated in ICU] (Figure 2(A)).

**Figure 2.**
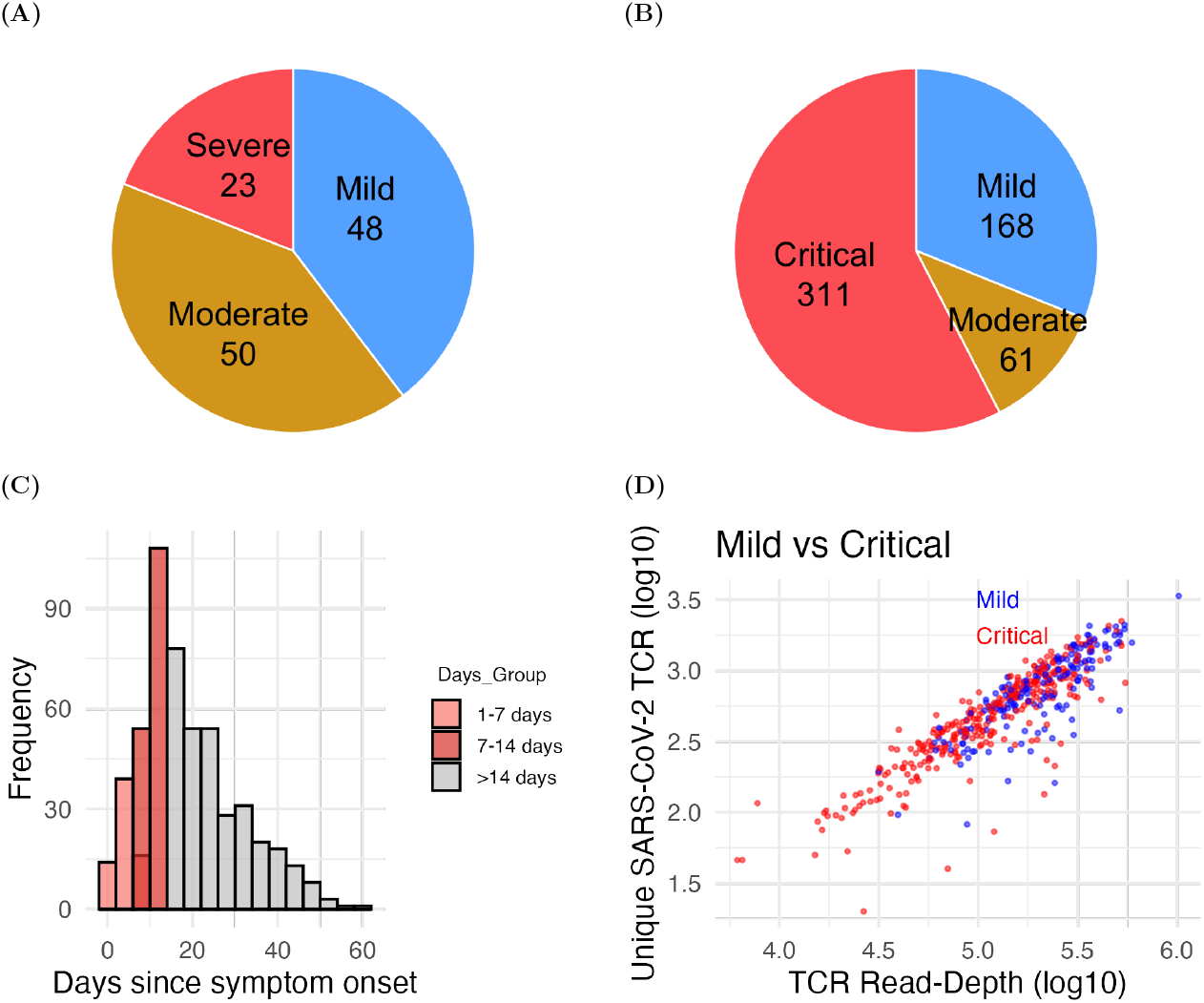
(A) The numbers of COVID-19 patients across severity groups from Su et al. dataset. (B) The numbers of COVID-19 patients across severity groups from Delmonte et al. dataset. (C-D) Additional summary of Delmonte dataset: the distribution of days since symptom onset (C) and the distribution of TCR read-depth versus richness (D). See Supplementary Figure 3 for the distribution of TCR read-depth versus richness involving moderate patients.

The dataset includes both single-cell and bulk-tissue TCR data. For the bulk TCR dataset, the average sequencing depth is around 10 million per individual, whereas for the single-cell dataset, it is approximately 10,000—nearly 0.1% of the bulk TCR dataset. Due to this lower coverage in single-cell data, most TCR clones are represented by only one or a few TCRs, making the richness and abundance metrics nearly identical. In comparison, the higher sequencing depth of the bulk TCR data allows for a clearer distinction between richness and abundance.

When estimating richness and abundance of SC2-TCRs, the approaches are similar for single-cell and bulk TCR datasets. However, single cell data provide the cell type information (i.e., CD4+ T vs. CD8+ T) for each TCR, enabling separate estimation for CD4+ and CD8+ TCRs. For example, when evaluating the abundance or richness of CD4+ T SC2-TCRs, we consider only TCRs derived from CD4+ T cells. In contrast, bulk TCR data lack CD4+/CD8+ cell-type labels, so richness and abundance estimates are based on all TCRs collectively. For example,

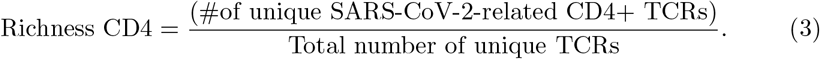

### Bulk TCR data from Delmonte et al

Delmonte et al. [14] collected TCR data from 677 blood samples of 540 COVID-19 patients, who were admitted to five hospitals in Italy between February 25, 2020 and September 9, 2020. These patients consist of 346 males and 194 females, with an median age of approximately 64 years. Among these 540 patients, 34, 196, and 310 were classified as mild, moderate to severe, and critical, respectively. The severity was ascertained based on the Diagnosis and Treatment Protocol for Novel Coronavirus Pneumonia (trial version 7) from the National Health Commission and State Administration of Traditional Chinese Medicine on March 3, 2020 [20].

The patients classified in the moderate to severe category exhibited varying levels of disease severity. To better quantify severity and to create more balanced comparison groups, we reclassified moderate to severe patients who were not admitted to the ICU and survived as mild cases. Using this reclassification, the updated distribution comprised 168 mild patients, 61 moderate to severe patients, and 311 critical patients (Figure 2(B)). Note that these severity categories are not directly comparable with those from Su et al. and thus we have analyzed the two datasets separately.

The samples were collected between day 1 and 60 post symptom onset (Figure 2(C)), with 43% of the cases sampled within 14 days, aligning with the early time point in Su et al [13]. Across the 677 samples, the average number of unique TCRs per sample was 58,884, with a range spanning approximately 3,000 to 300,000 (Figure 2(D), Supplementary Fig 3 (A)(B)).

## Results

### Results from Su et al. dataset

Su et al. [13] collected TCRs at two time points: an early time point (within two weeks of symptom onset) and a late time point (2-3 months post-symptom onset). The late time point was primarily used for studying long COVID. In this study, we focused on the TCRs collected at the early time point.

The single cell TCR data from Su et al. [13] provide precise differentiation between CD8+ TCRs and CD4+ TCRs. However, because of the low sequencing depth, most clones consist of only a single TCR, making it difficult to distinguish between TCR richness and abundance. Consequently, our analysis centers on TCR richness. Furthermore, we incorporated age as a covariate to further explore the relationship between SC2-TCR richness and the age of the patients.

The total number of CD4 + SC2-TCR from the MIRA set was limited (only 4. 3% of all MIRA TCRs), so in many samples, we did not observe any CD4+ SC2-TCRs. No significant difference was observed in the abundance of CD4+ or CD8+ SC2-TCRs across severity groups (Supplementary Figure 1(A,C)). There was a hint of a positive association between age and CD4+ SC2-TCRs in the severe group (Supplementary Figure 1(D)), though the data were too sparse to draw definite conclusions. For CD8+ SC2-TCR, there were trends of negative associations with age in all severity groups. However, these associations were not statistically significant, likely due to the sparsity of the data (Supplementary Figure 1(B)).

Next, using the bulk TCR data from Su et al. [13], we studied the associations between the richness and abundance of SC2-TCRs and disease severity. Severe disease was associated with lower abundance (Figure 3A) but higher richness (Supplementary Figure 2(A)). Since we expect weaker T cell response in early time point is associated with severe disease, these findings suggest that abundance, rather than richness, may serve as a more meaningful metric for quantifying the T cell response. The association between TCR abundance and disease severity was more pronounced when focusing on CD8+ SC2-TCRs (Figure 3B, p-value 0.03), while no significant association was observed for CD4+ SC2-TCRs (Figure 3C). Our analysis also revealed a negative relationship between age and the abundance of SC2-TCRs (Fig. 3D-F).

**Figure 3.**
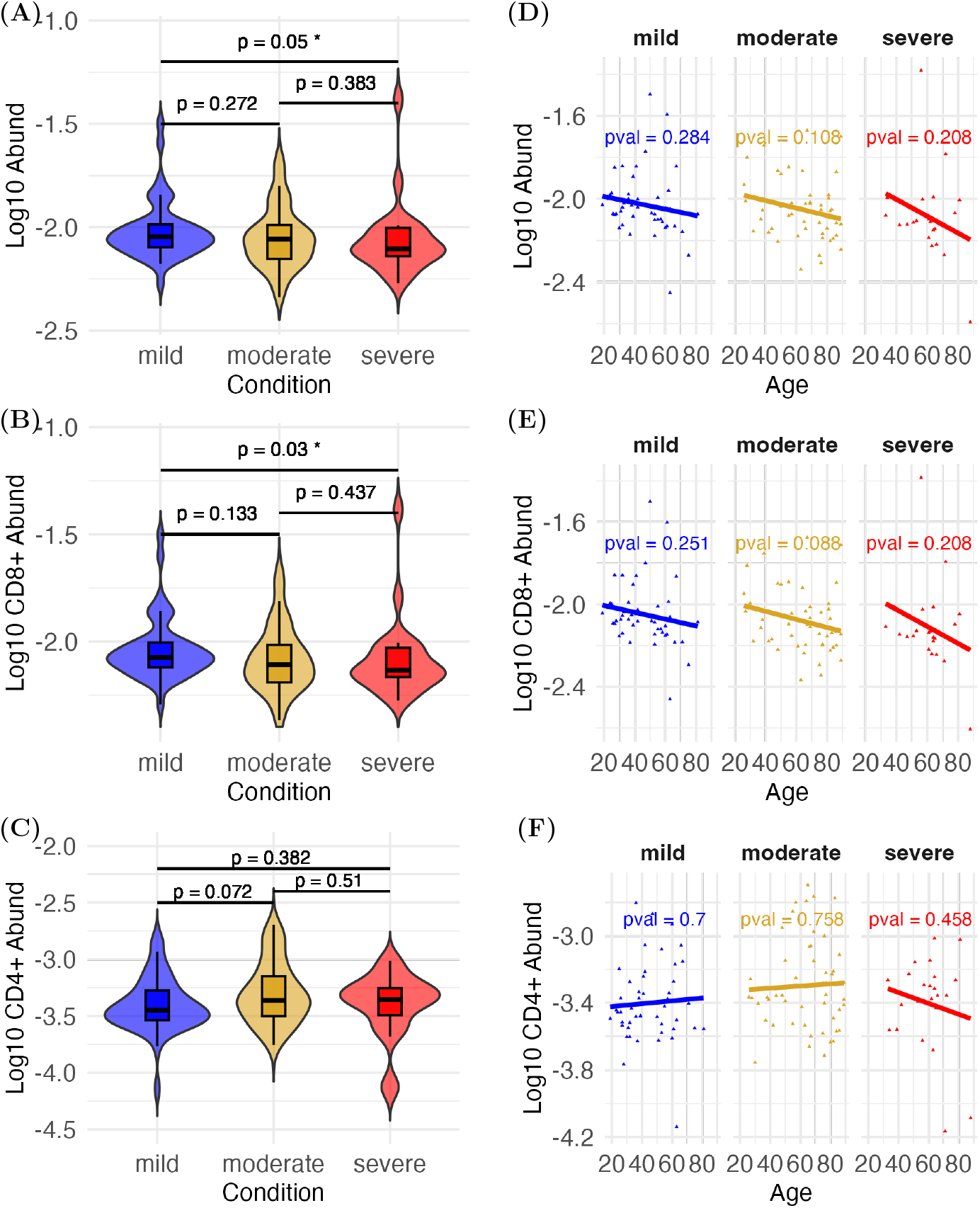
(A,B,C) Abundance of SC2-TCRs (ALL TCRs, CD8+ TCRs, CD4+ TCRs) in three severity groups. (D,E,F) Association between SC2-TCRs (ALL, CD8+ TCRs, CD4+ TCRs) and age.

We also evaluated the potential of using the abundance of SC2-TCRs to predict severe COVID-19. The prediction accuracy was moderate, with an AUC around 0.66 (Supplementary 6B), which might be partly due to the time frame in which the TCR samples were collected. As demonstrated in the next section in the analysis of Delmonte data, the association between TCR abundance and COVID-19 severity varies depending on the timing of TCR collection.

### Result from Delmonte et al. dataset

In the Delmonte et al. study [14], TCR data were collected at various time points up to 60 days post-symptom onset. To align with the time frame used in the Su et al. dataset, we initially focused on samples collected within 14 days of symptom onset. Our analysis revealed that the richness and abundance of SC2-TCRs could not effectively differentiate between mild and critical cases (Supplementary Figure 4(A), 5(A)). However, in critical cases, a negative correlation was observed between age and the abundance/richness of SC2-TCRs (Supplementary Figure 4(B), 5(B)), reinforcing the findings from Su et al. dataset.

We then extended the analysis to samples collected from day 1 to any time point up to day 30. A distinct pattern emerged when using all SC2-TCRs as the predictor. The ability to differentiate between mild and critical cases improved initially, peaking on day 8, and then gradually declined after this point (Figure 4A). Interestingly, when CD8+ and CD4+ TCRs were analyzed separately, their dynamics differed: CD8+ TCRs were more strongly associated with disease severity during earlier time points (before day 20), while CD4+ TCRs were associated with disease severity primarily at later time points (after day 20) (Figure 4A).

**Figure 4.**
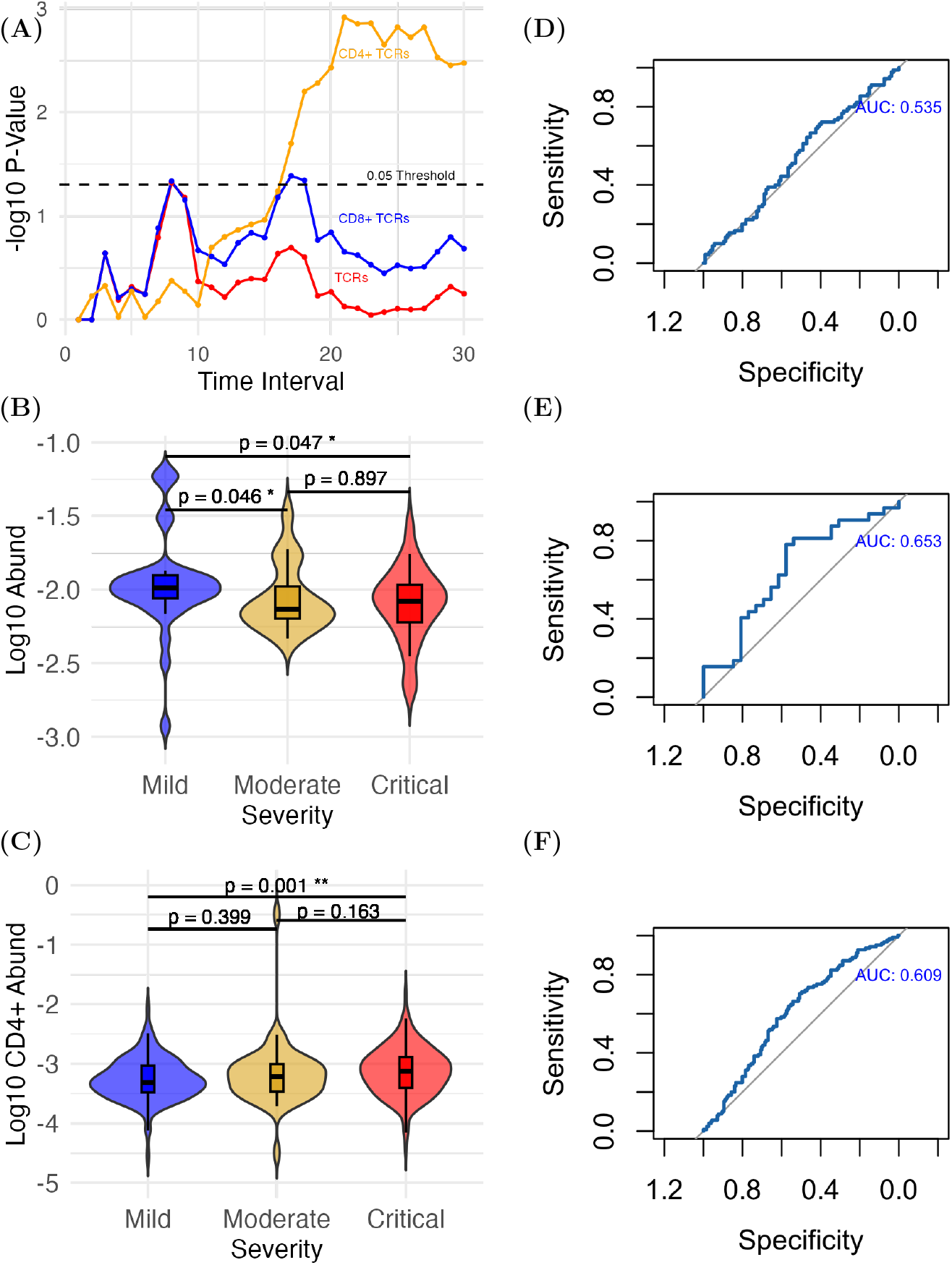
(A) Wilcox rank-sum test to compare the abundance of SC2-TCRs between mild and critical COVID-19 patients, when collecting TCRs up to a time point indicated by the x-axis. (B) Abundance of all SC2-TCRs in three severity groups within 8 days. (C) Abundance of CD4+ SC2-TCRs in three severity groups within 21 days. (D,E,F) ROC curves when classifying mild vs. critical patients while using (ALL, ALL, CD4+) SC2-TCRs using data collected by day (14, 8, 21) post symptom onset.

To explore these trends further, we selected two key time points for detailed analysis. Day 8 was chosen as the time when the abundance of all SC2-TCRs showed the strongest association with disease severity, and day 21 was selected for the significant association observed with CD4+ SC2-TCR abundance.

On day 8, the moderate and critical groups exhibited similar levels of SC2-TCRs, while the mild group showed significantly higher abundance (Figure 4 (B)). This similarity between moderate and critical groups may be explained by the reclassification of moderate cases discussed in the Methods section, as moderate cases included patients with severe disease (e.g., those admitted to the ICU). By day 21, CD4+ TCRs abundance was significantly higher in the critical group compared to the mild group (Figure 4 (C)).

To assess the ability of TCR abundance to distinguish between mild and critical patients, we performed AUC-ROC analysis. The results aligned with the findings from the Wilcoxon rank-sum test. On day 14, using all SC2-TCRs yielded an AUC of 0.535, indicating limited discriminatory power (Figure 4 (D)). However, on day 8 (Figure 4 (E)), the predictive performance was much better, with an AUC of 0.653 when using all SC2-TCRs to differentiate mild from critical cases. On day 21, using CD4+ TCRs as predictors yielded an AUC of 0.609, indicating moderate discrimination (Figure 4 (F)). When examining the AUCs across time points, the trends were similar to those of p-values: when using all SC2-TCRs, the AUC was higher in earlier time points. In contrast, when using CD4+ SC2-TCRs, the AUC was higher in later time points (Supplementary Figure 7).

### Prediction using top 1,000 TCRs with highest population frequencies

Sequencing millions of TCRs per patient could be cost-prohibitive for larger-scale studies. To address this challenge, we investigated whether a subset of public TCRs—those shared across patients—could predict disease severity. If successful, this approach could enable the development of targeted assays focusing on a smaller subset of TCRs.

Towards this end, we selected the top 1,000 TCRs with the highest population frequency and examined their association of COVID-19 severeness. The top 1,000 most popular TCRs from Delmonte et al. and Su et al. are highly consistent (> 97% overlap). Given the larger sample size and broader time range in the Delmonte data, which allows for the examination of dynamic patterns, we focused our analysis on this dataset. On average, each of these top TCRs was present in 25% of individuals, corresponding to approximately 135 out of the 540 individuals in the Delmonte dataset.

Next, we assessed the overlap of these 1,000 TCRs with SC2-TCRs. Among these top 1,000 TCRs, 30% overlap with MIRA TCRs, including 284 CD8+ TCRs and 18 CD4+ TCRs. This is particularly notable given that only 0.12% of all TCR clones in the Delmonte dataset overlap with MIRA TCRs. This substantial enrichment suggests that the top 1,000 TCRs are more likely to be associated with SARS-CoV-2 than the remaining TCRs.

We then repeated our association analysis using this subset of 1,000 TCRs, focusing on samples collected from day 1 to day 30 post-symptom onset. The results mirrored the patterns observed in the analysis using all TCRs (Figure 5): severe COVID-19 patients tend to have lower abundance of CD8+ SC2-TCRs at earlier time points, while higher abundance of CD4+ SC2-TCRs in later time points.

**Figure 5.**
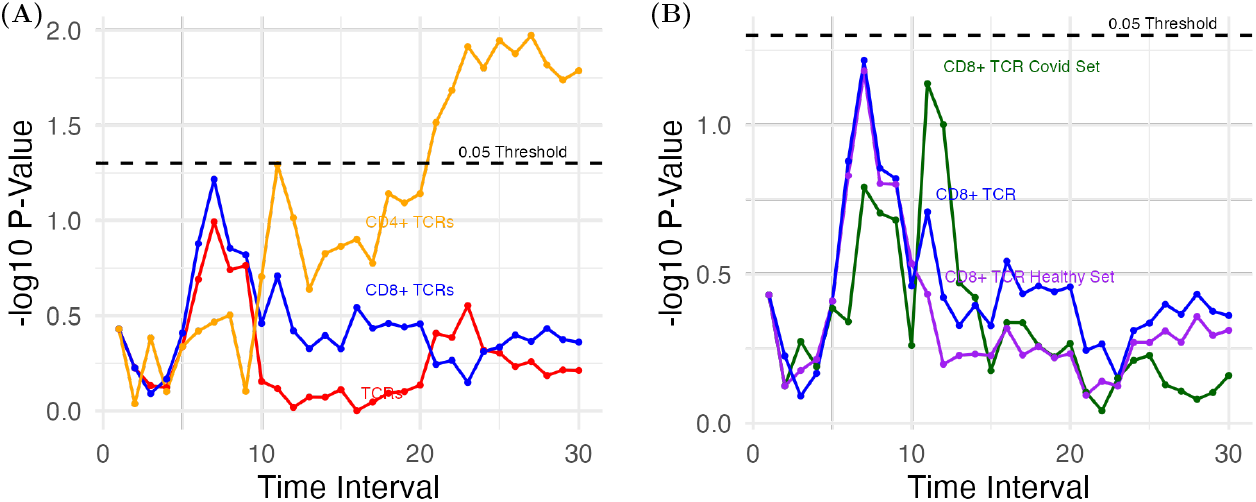
P-values for the association between COVID-19 severity and different sets of SC2-TCRs from the MIRA data, among the top 1,000 TCRs with highest population frequencies in the Delmonte data. (A) -log10(p-value) for three sets of TCRs: all TCRs, CD4+ TCRs and CD8+ TCRs. (B) Results for all CD8+ TCRs and two subsets: CD8+ TCRs from MIRA subjects who have SARS-CoV-2 exposure (CD8+ TCR Covid Set) or not (CD8+ Healthy TCR).

## Conclusions and Discussions

We investigated the dynamic associations between the abundance of SC2-TCRs and the severity of COVID-19. Our findings indicate that mild patients tend to have a higher abundance of SC2-TCRs (combining CD8+ and CD4+ TCRs) within the first 7-9 days of symptom onset, and the signal is mainly driven by CD8+ TCRs. After two weeks of symptom onset, critical patients show a significantly higher abundance of CD4+ SC2-TCRs. These findings highlight the importance of both timing and specific TCR subsets in understanding disease progression.

The observed patterns for SC2-TCRs align with expectations. During the early stages of infection, an elevated abundance of SC2-TCRs, reflecting a strong immune response, is associated with milder disease outcomes. However, as the infection resolves and recovery begins, the immune system down-regulates its activity, leading to the apoptosis of many virus-specific T cells. This reduction in T cell abundance may weaken the association between SC2-TCRs and disease severity over time.

Interestingly, the higher abundance of CD4+ SC2-TCRs at later time points (16 days+) is associated with more severe diseases. A possible explanation is that some CD4+ T cells may acquire cytotoxic functions during later stages of infection [13], potentially contributing to tissue damage. However, the relatively small number of CD4+ SC2-TCRs limits the ability to draw definitive conclusions.

Our analysis have a few limitations. First. due the limited sample size, we could not effectively study the associations between SC2-TCRs and disease severity at each time point. Instead, we used cumulative samples up to a time point, which would dilute the time precision. For example, while we found strongest association between CD4+ TCRs and disease severity around day 20, a day-to-day analysis may locate an earlier day of peak association. Second, while we found significant associations between TCR metrics and disease severity, the associations were not strong enough so that the TCR metrics could be an accurate predictor. A likely reason is that the MIRA TCR dataset used in this paper may include false positive SC2-TCRs and may miss some SC2-TCRs as well. In addition, it only have TCR beta-chains. We expect that given a more comprehensive and accurate collection of SC2-TCRs, particular with both alpha and beta chains, our method can be used to generate a more accurate predictor of COVID-19 severeness.

## Data Availability

The MIRA data used in this work can be accessed through the following link: A large-scale database of T-cell receptor beta (TCRb) sequences and binding associations from natural and synthetic exposure to SARS-CoV-2 (https://clients.adaptivebiotech.com/pub/covid-2020; DOI: 10.21417/ ADPT2020COVID).

The Delmonte data used in this work can be accessed through the following link: Perturbations of the T-cell receptor repertoire in response to SARS-CoV-2 in immunocompetent and immunocompromised individuals (https://clients.adaptivebiotech.com/pub/delmonte-2023-jaci; DOI: https://doi.org/10.21417/OMD2023JACI).

The single-cell TCR data from Su et al. 2020 study [13] can be accessed through Array Express under the accession number: E-MTAB-9357.

## Code Availability

The code used in this study is available on GitHub at https://github.com/Sun-lab/TCR_prediction/tree/main. The repository contains all the scripts and instructions required to reproduce the analyses and results presented in this paper.

## Notes

### Competing Interest Statement

J.R.H. is founder and board member of PACT Pharma. J.R.H. is a board member of Isoplexis.
J.D.G. declared contracted research with Gilead, Lilly, and Regeneron.

