## Supplementary Materials for "Association between COVID-19 Disease Severity and T Cell Receptor Repertoire"

November 26, 2024

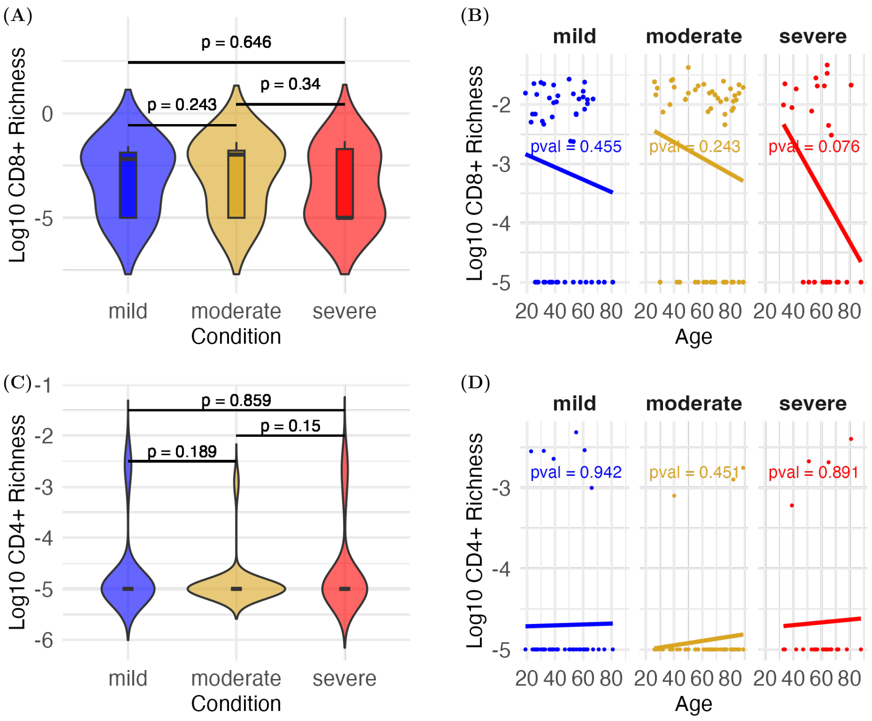


Figure 1: Analysis of SARS-CoV-2 associated TCRs (SC2-TCRs) in Su et al. [1] single cell TCR data. (A, C) Comparison of the richness of CD4+/CD8+ SC2-TCRs in three severity groups. The statistical significance of the difference between any two groups is assessed by Wilcox-Rank-Sum test. (B, D) Association between CD4+/CD8+ SC2-TCRs and age. The significance of the associations is assessed by linear regression.

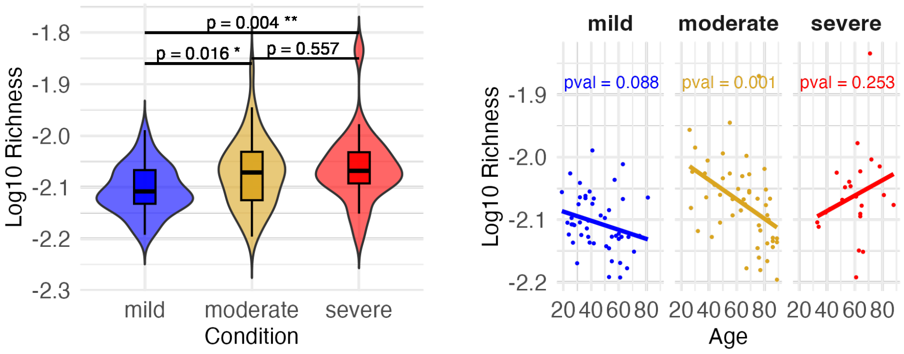


(A) (B)

Figure 2: Analysis of TCR richness using the bulk TCR data from Su et al. [1]. (A) Comparison of the richness of SC2-TCRs across three severity groups. (B) Association between the richness of SC2-TCRs and age.

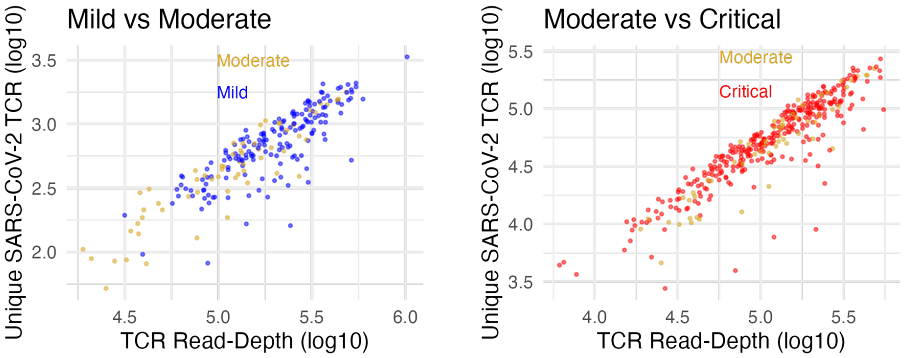


(A) (B)

Figure 3: Analysis of TCR richness using bulk TCR data from Delmonte et al. (2024) [2] (A) Comparison of SC2-TCR richness (measured by the total number of SC2-TCRs on y-axis) between the mild and moderate groups. (B) Comparison of SC2-TCR richness between the moderate and critical groups.

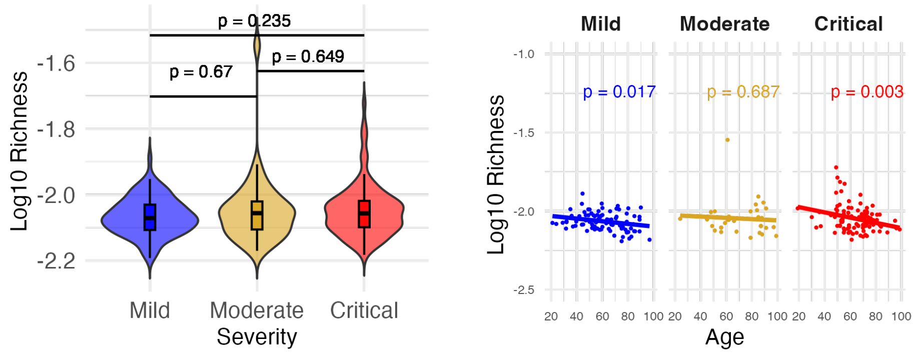


(A) (B)

Figure 4: Analysis of TCR richness using the part of the bulk TCR data from Delmonte dataset [2] that were collected within 14 days of symptom onset. (A) Comparison of SC2-TCR richness across three severity groups. (B) Association between SC2-TCR richness and age.

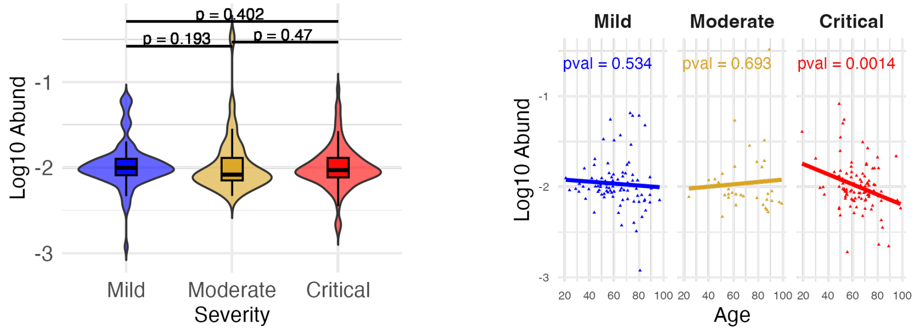


(A) (B)

Figure 5: Analysis of SC2-TCRs (collected within 14 days of symptom onset) in Delmonte dataset. (a) Abundance of SC2-TCRs across severity groups. (b) Relationship between SC2-TCR abundance and age across severity groups.

P values from Wilcox RankSum Test for Abundance of SARS_CoV_2-Related TCR in Mild vs Severe

| DATA Type | MIRA TCR sets | P Value |
|---|---|---|
| Bulk | ALL TCRs | 0.049 |
| Bulk | CD8+ TCRs | 0.03 |
| Bulk | CD4+ TCRs | 0.382 |
| Single Cell | ALL TCRs | 0.441 |
| SC CD8 | CD8+ TCRs | 0.61 |
| SC CD4 | CD4+ TCRs | 0.859 |

(A)

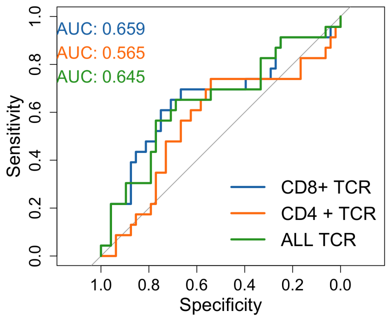


(B)

Figure 6: Summary of the associations between the abundance of SC2-TCRs and COVID-19 severeness using Su et al. data. (A) Wilcox rank-sum test p-values for comparisons using different types of TCR data and different MIRA TCR sets. (B) ROC curves for three bulk TCR predictors.

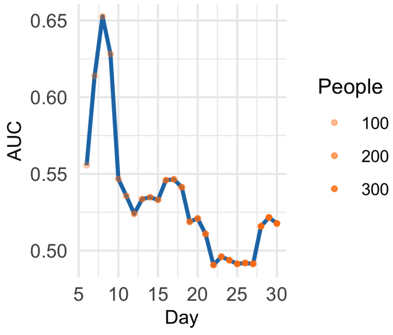


(A)

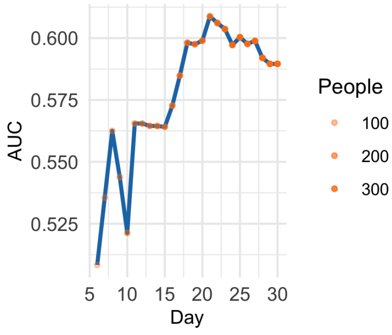


(B)

Figure 7: The AUC when classifying mild versus critical patients using the abundance of all SC2-TCRs (A) or CD4+ SC2-TCRs.(B)
